# Effects of DCM leaf Extract of *Gnidia glauca* (Fresen) on Locomotor Activity, Anxiety and Exploration-Like Behaviors in High Fat Diet-Induced Obese Rats

**DOI:** 10.1101/786103

**Authors:** Wycliffe Makori Arika, Cromwell Mwiti Kibiti, Joan Murugi Njagi, Mathew Piero Ngugi

**Affiliations:** Department of Biochemistry, Microbiology and Biotechnology, School of Pure and Applied Sciences, Kenyatta University, P.O. Box 43844-00100, Nairobi, Kenya; Department of Pure and Applied Sciences, Technical University of Mombasa, P.O. Box 90420 - 80100, Mombasa Kenya; Department of Environmental and Occupational Health, School of Environmental Sciences, Kenyatta University, P.O. Box 43844-00100, Nairobi, Kenya

**Author notes:** **Corresponding Author:** Wycliffe Makori Arika Department of Biochemistry Microbiology and Biotechnology Kenyatta University, P.O. Box 43844-00100, Nairobi, Kenya /+254722873150.

**Keywords:** Obesity, Anxiety, Locomotor activity, Exploration, high fat diet, *G. glauca*, Open Field Arena

## Abstract

Obesity is the main component of metabolic syndromes involving distinct etiologies which target different underlying behavioral and physiological functions within the brain structures and neuronal circuits. An alteration in the neuronal circuitry stemming from abdominal or central obesity stimulates a cascade of changes in neurochemical signaling that directly or indirectly mediate spontaneously emitted behaviors such as locomotor activity patterns, anxiety and exploration. Pharmacological agents available for the treatment of neurologic disorders have been associated with limited potency and intolerable adverse effects. These have necessitated the upsurge in utilization of herbal prescriptions due to their affordability, easy accessibility and are firmly embedded within wider belief systems of many people. *Gnidia glauca* has been used in management of many ailments including obesity and associated symptomatic complications. However, its upsurge in use has not been accompanied with authentication of these folkloric claims. The present study aimed to determine the modulatory effect of Dichloromethane leaf extract of *Gnidia glauca* on locomotor activity, exploration and anxiety-like behaviors in high fat diet-induced obese rats in an Open Field Arena. Obesity was experimentally induced by feeding the rats with prepared high fat diet and water *ad libitum* for a period of 6 weeks. The *in-vivo* anti-obesity effects were determined by oral administration of *G. glauca* at dosage levels of 200, 250 and 300mg/kg body weight in high fat diet-induced obese rats from the 6^th^ to 12^th^ week. Phytochemical analysis was done using gas chromatography linked to mass spectroscopy. Results indicated that *Gnidia glauca* showed anxiolytic effects and significantly increased spontaneous locomotor activity and exploration-like behaviors in HFD-induced obese rats. The plant extract also contained phytocompounds that have been associated with amelioration of the main neurodegenerative mediators viz inflammation and oxidative stress. These findings provide ‘qualified leads’ for the synthesis of new alternative therapeutic agent for management of neurologic disorders. However, there’s need to conduct toxicity studies of *Gnidia glauca* to establish its safety profiles.

## 1.1 Introduction

Metabolic abnormalities stemming from abdominal or central obesity are increasingly linked to impairments in central nervous system (CNS) function [1]. The hypertrophied and hyperplastic adipose tissue stimulates a cascade of changes in neurochemical signaling that directly or indirectly mediate behaviors [2,3]. Ostensibly, this degree of relative adiposity is implicated in a wide range of neuro-behaviors among which are, spontaneously emitted behaviors (activity patterns, anxiety and exploration), motivated behaviors (feeding, drinking, sexual behavior) and operant performance, attentional processes, learning and memory [4,5].

The obese phenotype and/or chronic exposures to high-fat diet (HFD) markedly exacerbates the odds of developing spontaneously emitted behaviors [6]. A study indicated that mice fed on HFD for 12 weeks showed reduced locomotor and exploratory behaviors in open field tests and elevated plus maze as well as depressive-like features characterized by reduced ambulatory activity in the forced swim tasks [5].

Exploration is one of the main domains of behaviors referring to the tendency to investigate a novel environment [7]. It is closely related to curiosity [7]. The cognitive map theory postulates novelty as misrepresentation of an item or place in the cognitive mapping/locale system [8]. The locale system is located within the hippocampus containing mental representations of previously perceived stimuli. Therefore, the hippocampal system supposedly signals a lack of information about the current environment and exploration becomes a direct response to the mismatch detected [8].

Chronic exposure to obesogenic diets is often associated with physical inactivity due to altered coordination of motor and reflexive responses [9,10]. Increased adiposity alters motor function through enhanced decrements in balance, muscle strength, and coordination [11]. Deficits in motor performance might be due to alterations in the striatal dopaminergic signaling or in the cerebellum [12,13]. Deficiencies in dopamine synthesis, striatal dopamine release, as well as defective striatal dopamine receptors are associated with impairments in striatal dopamine function [14,15].

Anxiety disorders, being the most prevalent mental disorders, globally contribute to reduced quality of life and predisposes affected individuals to other psychiatric comorbidities [16,17]. Anxiety, fear, chronic worry, muscle tension, panic attacks and apprehension are the main psychological symptoms while physical symptoms involve chest dysphoria, fatigue and tension [18]. Anxiety disorders are majorly categorized into specific and social phobias, panic disorder, obsessive-compulsive disorder, post-traumatic stress disorder and generalized anxiety disorder [18].

Studies have reported a positive association between obesity and anxiety disorders such as panic disorder, specific phobia and social phobia [19–21]. Obesity as a causal factor for anxiety involves several paths such as social discrimination against obese persons [22], low self-esteem in unfriendly social network [23], distress from illness burden (especially diabetes mellitus, asthma and cardiovascular diseases) and adverse drug effects [24]. Anxiety disorders as the causal factor for obesity has been correlated with disruption of hypothalamic-pituitary-adrenal (HPA) axis which results in dysregulation of autonomic functions [25,26]. These factors create another vicious cycle (obesity-anxiety cycle).

Inflammation and oxidative stress due to increased adiposity plays a pivotal role in the pathogenesis of neuropsychiatric disorders through their effect on the hypothalamus, amygdala and the hippocampus [27]. Chronic consumption of high fat diet stimulates production of proinflammatory cytokines (TNF-α) [28], interleukins (1β, 2, 6, 8 and 12) (Sahebkar, 2014), chemokines, immune cells, prostaglandins and nitric oxide which in turn precipitates hypothalamic-mediated oxidative stress [29]. The compromised redox homeostatic status characterized by attenuated antioxidant defenses activates brain’s resident immune cells, the microglia and astrocytes, to further produce inflammatory mediators, which exacerbates an oxidative damage in the hippocampus [30].

Pharmacological agents available for the treatment of neurologic and psychiatric disorders have had limited potency or intolerable adverse effects [31]. Therapeutic herbs and nutrients have, therefore, provided an effective alternative treatment with minimized side effects and capacity to potentiate the effect of prescribed medications [32]. The present study aimed to determine the modulatory effects of DCM leaf extract of *Gnidia glauca* on locomotor activity, exploration and anxiety-like behaviors in HFD-induced obese rats in an open field arena. The generated data will provide ‘qualified’ leads in drug design from this plant for the treatment and/or management of neurologic disorders.

## 1.2 Materials and Methods

### 1.2.1 Processing and Extraction of the Plant Material

Fresh leaves of *G. glauca* were dried on a shade at room temperature for 21 days. By use of an electric mill, the dried leaves of *G. glauca* were ground into fine powder. The milled plant sample was then kept at room temperature free from direct sunlight in a dry airtight plastic container prior to extraction. In one litre of dichloromethane (DCM), 500 grams of powdered sample of *G. glauca* was put and soaked for forty-eight hours. The dissolved compounds were decanted and filtered using muslin cloth into a dry clean conical flask. The filtrate was concentrated under reduced pressure by use of a rotary evaporator at 40°C to attain a semi-solid residue [33]. The yield of the plant extract was determined and subsequently refrigerated at −20° C prior to its analysis.

### 1.2.2 Obesity induction

Obesity was induced by feeding the experimental animals with a high-fat diet and regular supply of water for a duration of twelve weeks. The ingredient of the experimental diet was done according to a formula as described by Srinivasan *et al*. [34] with minimal moderation as indicated in Table 1.1.

**Table 1.1:** High Fat Diet Composition

| Ingredients | Diet (g/kg) |
| --- | --- |
| Lard | 290 |
| Powdered NPDL | 375 |
| Casein | 265 |
| Vitamin and mineral mix | 60 |
| Cholesterol | 10 |
| Corn oil | 10 |
| DI Methionine | 03 |
| Sodium Chloride | 01 |
| Yeast powder | 01 |

The obesity index was defined by Lee index. The Lee index was calculated according to the formula described by Lee [35].

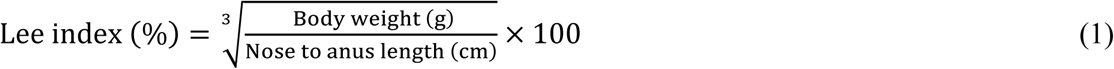

Rats with Lee obesity index value (equivalent to BMI≥30 in humans) of 310 and above were considered obese [35] and used in the study. Following exposure to HFD (except for normal control group) for 6 weeks, all the rats in the negative control, positive control and extract-administered experimental groups attained the target diagnostic value of obesity, indicating the end to the obesity induction phase. The Naso–Anal Length (NAL) (cm) of rats was measured by a non-extensible thread and readings taken using a ruler with an accuracy of 0.1 cm.

### 1.2.3 Experimental Design

Thirty female rats were randomly grouped into 6 different sets of 5 animals each. The normal control group (Group I) comprised of normal animals that were fed on a standard chow diet for 12 weeks. Group I animals never received any treatment. Group II (negative control) comprised of rats that were fed on HFD in a duration of 12 weeks to induce obesity. The positive control group (Group III) consisted of HFD-induced obese animals that received reference drug, Orlistat, from the 6^th^ to 12^th^ week. Group IV-VI (experimental groups) comprised of HFD-induced obese rats that were administered with the DCM leaf extract of *G. glauca* at different doses of 200, 250 and 300 mg/kg body weight from the 6^th^ to 12^th^ week. All the treated rats only fed on high-fat diet during the entire dosing period. Further, all the experimental rats received water *ad libitum* throughout the study period.

### 1.2.4 Open Field Arena

In order to determine whether *G. glauca* modulates gross locomotor activity, exploration and anxiety-like behaviors in HFD-induced obese rats, animals were tested in an open field arena [36] after 6 weeks of oral administration of therapeutic doses of the plant extract.

#### 1.2.4.1 Apparatus

The open field apparatus consisted of an open top box (72cm x 72cm) with 36 cm high walls. Blue visible lines were drawn on the floor using a marker into sixteen 18 x 18 cm squares (Figure 1.1). A center square (18 cm x 18 cm) was drawn in the middle of the arena (within the four inner squares) with a red marker (Figure 1.1). A 60-Watt white light bulb provided lighting. The floor was covered by a sheet of clear Plexiglas which was cleaned using 70% ethyl alcohol after testing each animal. Animal behaviors in the arena was recorded and tracked by an overhead video camera connected to a PC with Ethovision XT software (Figure 1.1).

**Figure 1.1:**
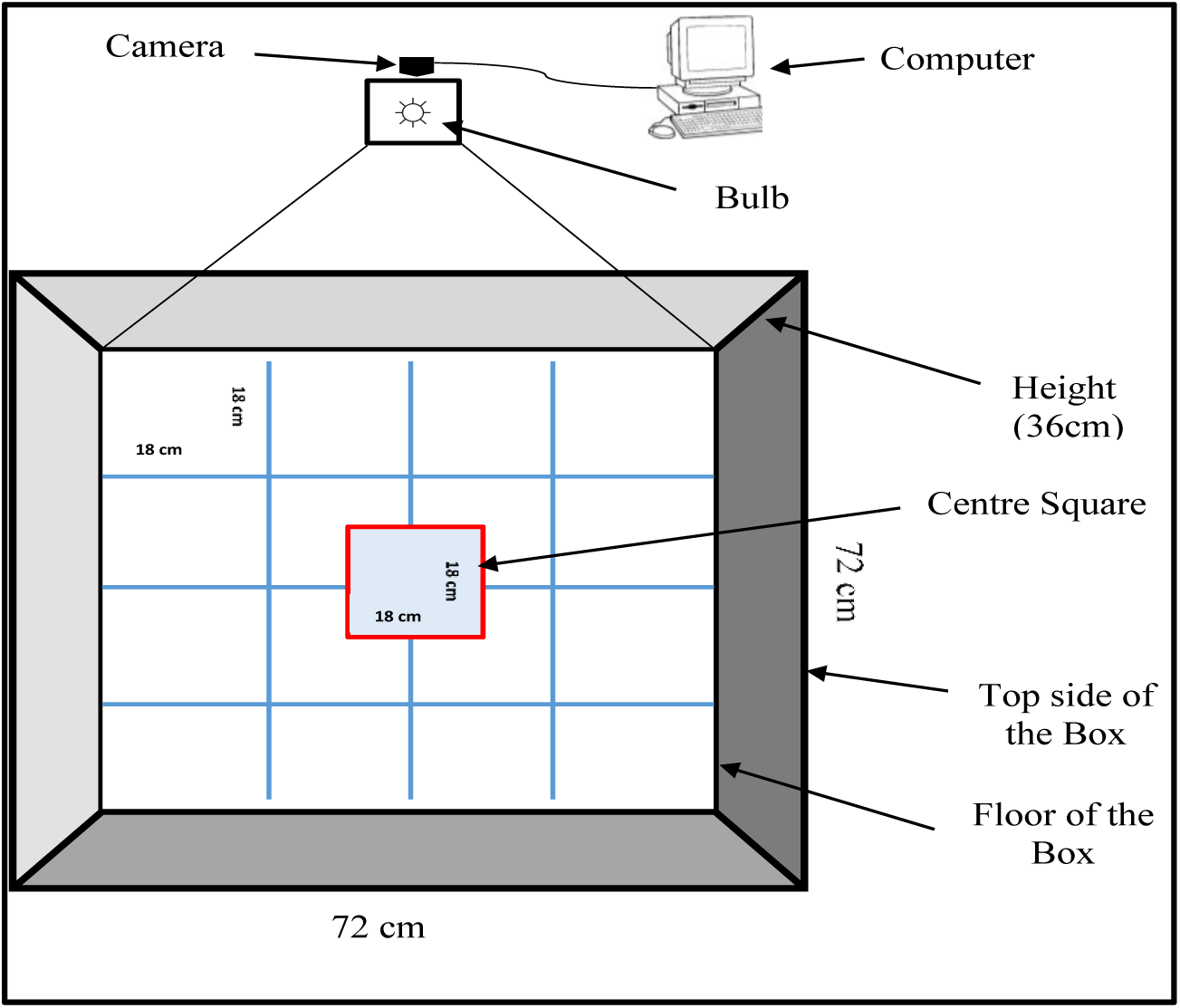
The Open Field Arena

#### 1.2.4.2 Experimental Procedure

The animals were transferred to the testing room in their home cages and allowed to acclimatize to this room prior to testing. Each rat was gently placed in the center of the open field arena and left freely to explore the arena for 5 minutes while recording scores of its behaviors. At the end of the 5-minute test period, animals were returned to their respective home cages. The Plexiglas was removed, cleaned and disinfected with 70% ethyl alcohol between each trial.

To determine the effect of the DCM leaf extract of *G. glauca* on locomotor activity, exploration and anxiety-like behaviors, the following animal behaviors were assessed: number of line/grid crossing (frequency with which the rats crossed one of the grid lines with all four paws), center square entries (frequency with which the rats crossed one of the red lines with all four paws into the central square), latency period (duration of stay in the central square), rearing (frequency with which the rats stood on their hind legs in the arena), stretch-attend-postures (frequency with which the rats demonstrated forward elongation of the head and shoulders followed by retraction to their original position), grooming (duration of time the animal spent licking or scratching itself while stationary), freezing (duration with which the animal was completely stationary), urination (number of puddles or streaks of urine) and defecation (number of fecal boli produced per animal) [36].

Locomotor activity for each animal was assessed by the sum of line/grid crosses and number of rears. The behavioral domains used to test exploration-like behaviors included the frequency of rearing, central square entries and latency in the central square. The animal behavioral domains used to assess anxiety were latency period, fecal boli score, frequency of urination, grooming, stretch attend postures and freezing.

### 1.2.5 Gas Chromatography-Mass Spectrometry Analysis

Sample analysis was determined using GC-MS (7890/5975 Agilent Technologies, Inc., Beijing, China) consisting of a gas-chromatograph integrated into a mass spectrometer instrument. The GC-MS was equipped with a HP-5 MS (5% phenyl methyl siloxane) low bleed capillary column of 0.25µm film thickness, 0.25mm diameter and length of 30m. An electron ionization system with ionization energy of 70Ev was used in GC-MS detection. Helium (99.99%) gas carrier was used at a consistent flow rate (1.25 ml/min) in split mode. The mass transfer line and injector temperature were set at 200°C and 250°C respectively. One microliter was used as an injection volume. Oven temperature was programmed from thirty-five degrees Celsius for five minutes, with an elevation of ten degrees Celsius per minute to two hundred and eighty degrees Celsius for 10.5 minutes, followed by fifty degrees Celsius per minute to two hundred and eighty-five degrees Celsius for 29.9 minutes with seventy minutes run time. The mass spectrometry operating parameters included: ionization energy, 70eV; ion source temperature, 230°C, relative detector gain mode, scan speed 1666 µ/sec, solvent cut time, 3.3 min; the interface temperature was 250°C, scan range 40-550 m/z.

### 1.2.6 Data Management and Statistical Analysis

To assess the performance in the open field arena, each determinant of the behavioral domain was recorded and tracked by an overhead video camera connected to a PC with Ethovision XT software. The data for each behavioral domain was exported to Microsoft® Excel spread sheet, where it was organized and later transferred to statistical software Minitab (Version 17.1) for analysis. The data was found to conform to the assumptions of parametric data. One-way ANOVA was used to test the significant differences among the normal control group rats, negative control group rats, Orlistat-treated group of rats and extract-treated group of rats. The data was further subjected to Tukey’s post hoc for pairwise comparison and separation of means. The criterion for significance was set at *p*≤0.01. The findings were presented in a table.

## 1.3 Results

### 1.3.1 Effect of DCM Leaf Extract of *Gnidia glauca* on Locomotor Activities, Anxiety and Exploration-Like Behaviors in HFD-Induced Obese Laboratory Rats

Treatment of HFD-induced obese rats with DCM leaf extract of *G. glauca* resulted in a significantly higher grid crossing score relative to the negative control group rats (*p*≤0.01; Table 1.2). Further, rats treated with the plant extract showed a higher grid crossing than those in the normal control group (*p*≤0.01). Administration of extract dosages of 250 and 300mg/kg body weight resulted in a significant increase in grid crossings than those of rats treated with the reference drug, Orlistat (*p*≤0.01; Table 1.2).

**Table 1.2:**
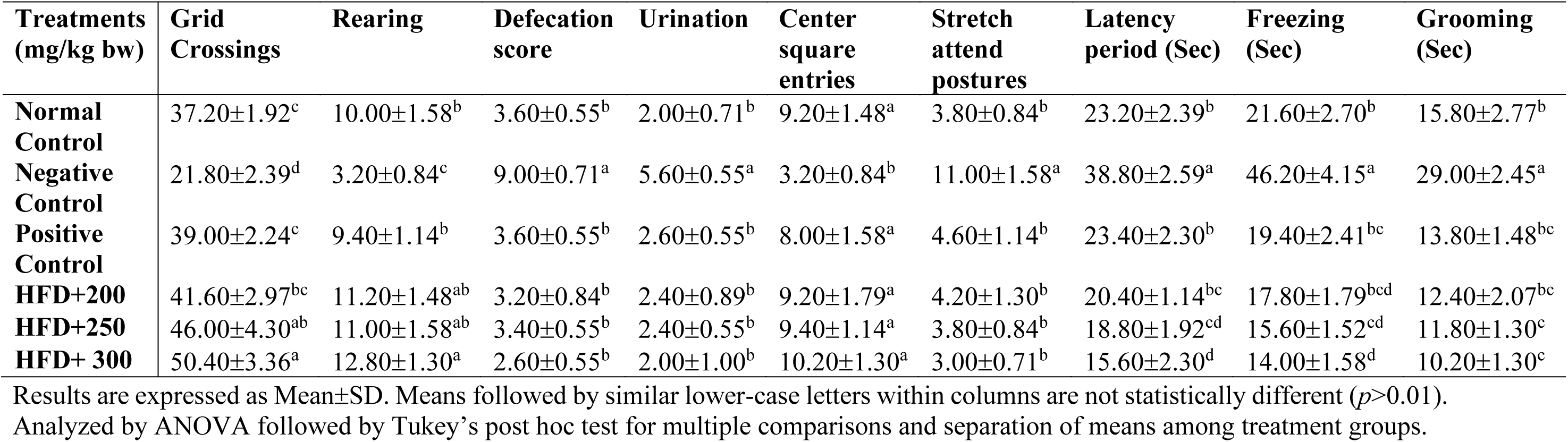
Effect of DCM Leaf Extract of *Gnidia glauca* on Locomotor Activities, Anxiety and Exploration-Like Behaviors In HFD-Induced Obese Rats

It was further observed that HFD-induced obese rats treated with the plant extract had a higher number of rearing episodes relative to HFD-fed untreated rats (Table 1.2). Besides, there was no significant variation in the number of rearing episodes observed among extract-treated rats at dosages of 200 and 250mg/kg body weight and those of normal control group rats and rats in the positive control groups (*p*>0.01). Results indicated higher defecation and urinating episodes in rats in the negative control group than those of rats in the extract-treated groups (*p*≤0.01). Similarly, the number of stretch attend postures and urinating episodes were significantly high in HFD-fed untreated rats relative to extract-treated group of rats (*p*≤0.01). Further, it was observed that the number of visits to the central square after the initial exit were more on those groups of rats that were treated with the plant extract and the reference drug, Orlistat, than HFD-fed untreated obese rats (Table 1.2). Remarkably, no significant difference in defecation, urination, center square entries and stretch attend postures were observed among rats in the extract-treated, positive control and normal control groups (*p*>0.01; Table 1.2).

The results also showed that the HFD-fed untreated obese rats in the negative control group had a significantly longer latency period in the central square upon entry into the open field arena than rats treated with *G. glauca* leaf extract (*p*≤0.01). Treatment of rats at a dosage levels of 250 and 300mg/kg body weight of the extract resulted in a shorter latency period in the central square than rats administered with the reference drug, Orlistat (Table 1.2). However, the latency period was statistically similar in rats treated with the reference drug, Orlistat and those in the normal control group (*p*>0.01).

The results also revealed that rats in the negative control group substantially froze longer upon exposure to the open field arena than rats treated with the three extract doses (Table 1.2). However, the effect was not statistically significant among rats treated with the reference drug, Orlistat and those treated with the plant extract at dosage levels of 200 and 250mg/kg body weight (*p*>0.01). Treatment of rats with the highest extract dose of 300mg/kg body weight significantly reduced immobility time (freezing period) than Orlistat-treated rats (*p*≤0.01; Table 1.2).

Results also demonstrated that administration of *G. glauca* leaf extract significantly reduced the grooming behavior in HFD-induced obese rats relative to untreated obese rats in the negative control group (*p*≤0.01). However, the propensity to groom in the extract-treated groups of rats and those treated with the reference drug, Orlistat, were comparable (*p*>0.01; Table 1.2).

### 1.3.2 The Concentrations of Compounds Identified in DCM Leaf Extract of *Gnidia glauca*

The gas chromatography-mass spectrometry analysis of DCM leaf extract of *G. glauca* indicated the presence of Oleic acid, γ-Sitosterol, Curcumin, Quercetin, Stilbenes, Phytol, Octadecanoic acid (Stearic Acid), Gallocatechin-catechin flavan Ferulic acid, Flavonols and others (Table 1.3).

**Table 1.3:**
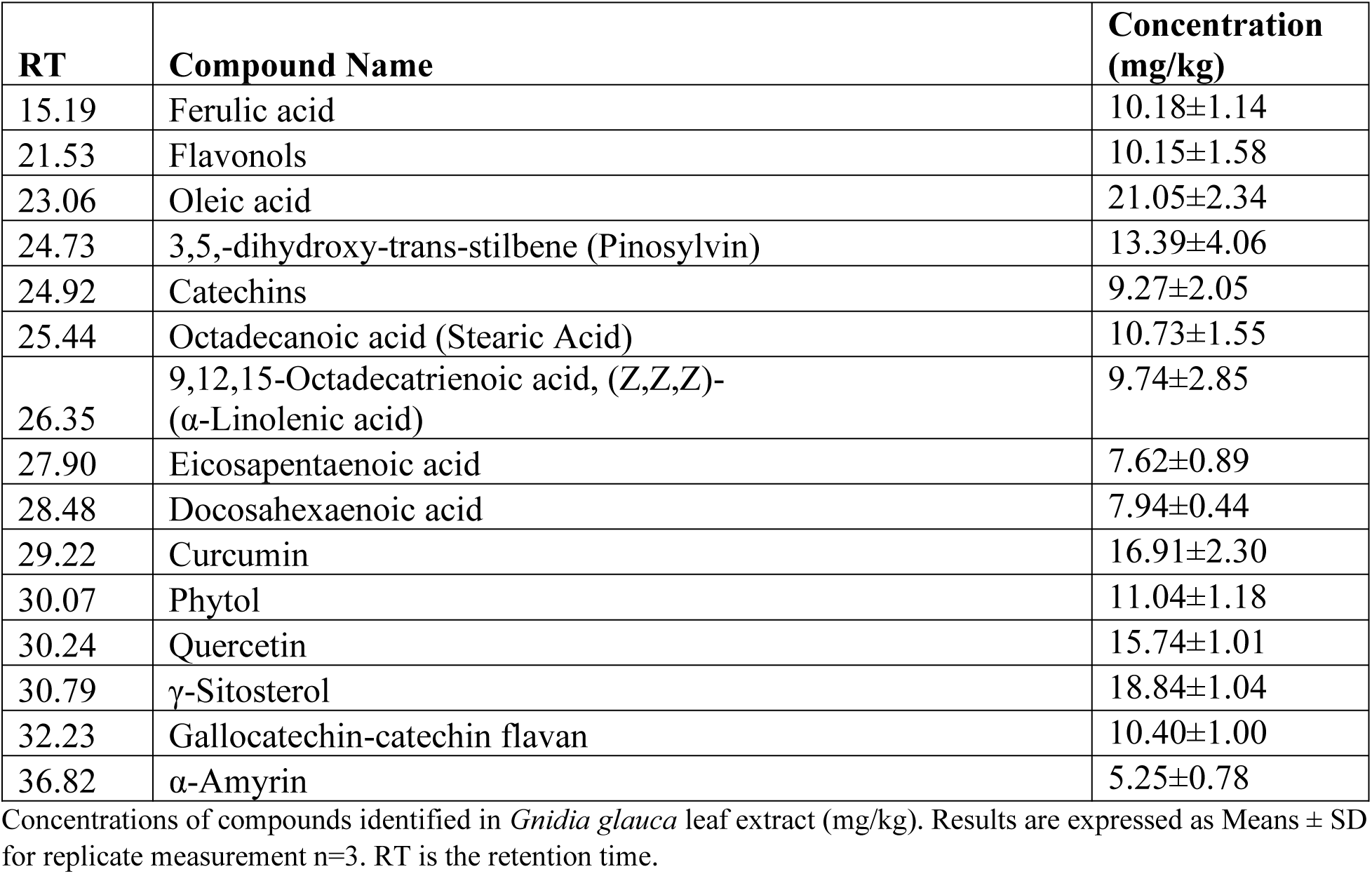
Quantity of Phytochemical Compounds in DCM Leaf Extract of *Gnidia glauca*.

## 1.4 Discussion

The open field test provides simultaneous measures of locomotion (ambulatory activity), exploration and anxiety (emotionality) [37]. It is a model not only useful in the assessment of behavioral performance of the test animals but also contributes knowledge on the neurobiological mechanisms mediating behaviors [37].

Locomotor/ambulatory activity is the function of performance on motor tasks while exploration or novelty may involve some quality never previously experienced or familiar items arranged in unfamiliar ways [38]. Exploratory behavior is thus curiosity and attraction to novelty [38]. Animal behaviors such as frequency of line crosses, frequency of rearing, central square entries and latency in the central square are used as measures of locomotor activity and exploration. A higher frequency of these parameters indicates increased locomotion and exploration and vice versa [37]. The present study indicated a number of clear effects on these behavioral domains.

The frequency of line/grid crosses measures the horizontal exploration or locomotor behavior and represents the horizontal distance covered [38]. Line crossing is the frequency with which the rat cross each of the lines with all four paws. In the present study, line crossing was significantly (*p≤*0.01) increased in the extract-treated rats as compared to the HFD-fed untreated obese rats. Similarly, administration of ethanolic extract and acetone extract of *Cedrus deodara* increased locomotor activity in neonatal rats [39]. In normal circumstances, rats naturally move in order to find location of feeds, gather nesting materials, search for nesting places and sexual partners or flee themselves from enemies [40]. However, the reduced locomotor activity observed in HFD-fed untreated obese rats could be due to the overweight nature of the animals, generalized muscle fatigue and/or increased behavioral despair [38]. Besides, the diminished ambulatory/locomotor activity could be as a result of the damage in the primary motor area and /or distress from illness burden associated with increased adiposity [25,38]. The ability of the extract to reduce body weight might be attributed to the observed increased ambulatory activity in extract-treated rats. Moreover, the extract might have resulted in a positive effect on striatal dopaminergic signaling through increased striatal receptor sensitivity and dopamine synthesis thereby improving motor activity [13].

The frequency of rearing or vertical exploration was significantly decreased in HFD-fed untreated obese rats as compared to their treated counterparts (*p≤*0.01). Rearing measures exploratory behavior or otherwise vertical locomotor activity. When rearing, the animal stands upright on its hind limbs often using their tail as support with its fore limbs freely suspended in the air or resting on the wall of the open field arena. Through rearing, olfactory signals can be taken in from the air, as well as visual cues [41]. Locomotor activity is driven by exploration since its reduced form could possibly reflect reduced exploration as it is accompanied by reduced rearing frequency [41]. Previous studies demonstrated that oral administration of ethanolic extracts of *Nauclea latifolia* and *Emilia sonchifolia* increased locomotion and exploratory activities as evidenced with a high frequency of rearing in mice [38].

The state of being obese is associated with decreases in the motor output, often termed as “physical inactivity” [10]. Chronic exposures to obesogenic diets contribute to striatum damage thereby affecting the dopamine synthesis and release as well as striatal-dopamine receptor function [13]. The striatal-dopamine plays a key role in the proper control of movement and, therefore, its impairment contribute to physical inactivity in obesity akin to classical movement disorders such as Parkinson’s disease [13,42]. The motivated locomotor and exploratory behaviors observed in extract treated rats could been linked to facilitated dopamine synthesis and release as well as restored striatal dopamine receptor function [43,44].

The chronic mobility problems of joints and muscles in obese patients is largely contributed by alteration of motor circuitry in the brain [13]. Besides, the obesity-induced adaptations due to altered motor circuitry could continue to contribute to physical inactivity even after weight loss [13]. The reduction of expression levels of brain-derived neurotrophic factor (BDNF) and its tyrosine kinase receptor, TrkB, in hypothalamic nuclei affects the strength of synaptic connections or dendritic spine density leading to altered satiety signals and locomotor activity [45]. High fat diets potentiate an oxidative attack on the brain resident cells resulting in an activation of cholinergic motor inhibitory system [46]. Alteration of the activity of acetylcholinesterase (AchE) and damage to the peripheral muscle due to necrosis of skeletal muscle fibres enhances reduction of locomotor activity in animal models [47].

The increased ambulatory or spontaneous physical activity (SPA) characteristic to extract-treated rats could also be as a result of the action of neuropeptide, orexin A, independent of feeding behavior [48]. Orexin A robustly stimulates spontaneous physical activity and non-exercise activity thermogenesis [49,50]. Central administration of orexin-A (into the hypothalamic paraventricular nucleus) was found to increase rearing frequency and locomotor activity in rats [50]. The orexin neurons project to the dopaminergic neurons in the substantia nigra that innervate the striatum and forms a critical component of motor activity [51]. Therefore, an alteration in the expression of orexin, and/or its signaling, could exacerbates spontaneous physical inactivity and contribute to weight gain [51].

The test for anxiety is usually based on the conflicting tendencies of rats to explore a novel environment in contrast to the aversive features of a brightly lit open arena or an elevated space [37]. Moreover, in the open field arena, this behavioral domain may be mediated by two key factors namely, agoraphobia and individual testing. Agoraphobia is a function for anxiety based on size of the test area relative to the size of an animal while individual testing is a function for anxiety based on separation of an animal from its social group [52]. Animal behaviors such as, increased latency period, greater fecal boli score, higher frequency of urination, increased grooming period, less rears, higher frequency of stretch attend postures and increased freezing duration are used as measures of anxiety. A higher frequency or an increased duration of these parameters indicates increased anxiety [37].

Analysis of stretch attend postures (SAP) revealed a significantly (*p≤*0.01) increased frequency in obese untreated rats relative to extract-treated rats. The SAP is the frequency with which the animal demonstrated forward elongation of head and shoulder followed by retraction to its original position. These are risk assessment behaviors of fear and anxiety which indicates that the animal is hesitant to move from its present position of comfort to a new position. Thus, decreased levels of this behavior are indicative of a low level of anxiety and fear and vice versa [53]. These results were consistent with the finding that mice with HFD-induced obesity demonstrated a high frequency of SAP relative to obese mice treated with the herbal extracts from *Morus alba*, *Melissa officinalis*, and *Artemisia capillaris* [54].

Results also showed that the extract-treated rats had an increased frequency of entry to the inner zone of the open field arena relative to HFD-fed untreated obese rats (*p≤*0.01). A high frequency of movements into the center of the arena in open field tests is reflective of reduced anxiety, increased locomotor activity and exploration [37]. Rats are generally thigmotactic, they avoid open areas and prefer moving alongside walls where they perceive tactile stimuli via their vibrissae [41]. However, when the animal is less anxious their exploratory behavior increases and tend to move all over the holding cage or arena. Consistent with the present study, previous studies observed that treatment of mice with hydroalcoholic extract of *Coriandrum sativum* increased frequency of entry to the inner zone of the open field arena [55]. The sedative and muscle relaxant effects of *Coriandrum sativum* are indicative of its anxiolytic effects [55].

The HFD-fed untreated group showed longer latency period in the central square upon entry into the open field arena, an indicator of higher anxious states due to anxiogenic effects of chronic exposure to high fat diets. The quicker the retreat from the center square of the arena in extract-treated rats is indicative of increased automatic and exploratory behaviors due to extracts anxiolytic effects [56]. These findings were in agreement with a study that demonstrated that HFD-induced obese rats showed less explorative interest due to lower cross lattice number and reduced percent of time spent in the center of the arena and open arms [57]. The reduction in explorative interest in HFD-induced obese untreated rats appear to be symptoms of depressive disorders consistent with those observed in patients suffering from anxiety disorders [58]. Current studies have demonstrated that chronic intake of HFD have led to depressive- and anxious-like behaviors [57,59].

Immobility time (freezing duration) was significantly (*p≤*0.01) increased in HFD-fed untreated rats compared to extract-treated rats, an indicator of increased anxious state and hypoactivity or impaired locomotor activity [37]. Freezing often occurs in response to a sudden change in the surroundings where the animal usually stands still with its forelegs raised while looking up. Previous studies demonstrated that HFD-induced obese rats exhibited a significantly low frequency of rearing as compared to rats treated with therapeutic doses of aqueous extract of *Ginkgo biloba* [60]. The anxiolytic effects of the *G. glauca* leaf extract might be accompanied by increases in the brain levels of monoamines such as serotonin, norepinephrine, and dopamine [61]. Serotonin and norepinephrine are neurotransmitters that plays a key role in mood regulation [62,63].

Increased fecal boli and urination scores observed in HFD-fed untreated obese rats are suggestive of fear and anxious states. Comparatively, the low scores of these parameters in the extract-treated rats maybe attributed to presence of bioactive chemicals responsible for down-regulation of receptors and connectivity in the amygdala, a key center of fear [25,37]. Previous studies have demonstrated that high-fat feeding and obesity increases the production of BDNF and phospho-CREB in the striatum contributing to negative emotional states and depressive-like symptoms [5]. This biochemical alterations in the brain reward circuitry could be implicated for the observed increased fecal boli and urination score in HFD-fed untreated obese rats.

Grooming is a de-arousing self-directed behavior associated with anxiety upon displacement of animals into a novel environment or aversive situations such as an open field arena [64]. Grooming duration is the time the animal spends licking or scratching itself with paws and face washing actions while in stationary position. This stereotypical behavioral sequence is usually increased in anxious states. Anxiolytic drugs, however, reduces the grooming behavior, whereas anxiogenic drugs facilitate grooming [41]. Treatment with therapeutic doses of the plant extract led to a significant (*p≤*0.01) decrease in grooming duration relative to HFD-fed untreated obese rats. Consistent with this study, the open field tests performed to test the neurobehavioral effect of *Nauclea latifolia* and *Emilia sonchifolia* [38] and *Mammea africana* [65] in rodents indicated reduced grooming frequency together with increased spontaneous locomotion and exploratory activities [66].

Self-grooming is a highly stereotyped patterns of sequential movements (syntactic chain pattern) that is modulated by circuits that incorporate the basal ganglia such as the striatum, substantia nigra and nucleus accumbens in the forebrain [64,67]. Striatal circuits majorly sub-serve the basal ganglia and are mainly involved in learning, motivation and motor sequencing [68]. Lesions of the striatum completely impairs sequential syntactic self-grooming chains [64]. The limbic circuitry that include the amygdala and the hypothalamus also modulates self-grooming behavior in rodents. The amygdala mainly modulates motivational states, such as desire, fear and anxiety [69]. Studies showed correlations between reduced dopamine release within the amygdala and increased anxiety-like behavior in low and high grooming rats respectively [70].

The hypothalamic paraventricular nucleus is another limbic region that has been implicated in the regulation of self-grooming in rodents [70]. Besides, the hypothalamic–pituitary stress-related hormones such as corticotropin-releasing hormone (CRH) and adrenocorticotropic hormone (ACTH) also influences self-grooming in rodents [64]. The anxiolytic effects observed in the extract-treated rats as evidenced by reduced self-grooming, increased exploratory and locomotor activities and could be attributed to the extracts effect on the dopamine release in the nigrostriatal and mesolimbic systems [71]. Dopamine plays a critical role in locomotor function, self-grooming and other complex behavioral patterns [64]. The *G. glauca* leaf extract might have also contributed to reduction of stress-induced self-grooming by enhancing the GABAergic tone through attenuating the intensity of the perception of anxiogenic stimuli [71,72].

The mechanistic basis underlying obesity as a causal factor for anxiety could relate to oxidative stress and inflammation [30]. Studies have demonstrated that exposures to chronically high energy diets influences the activity of glial cells that mediates endogenous immune system within the microenvironment in the CNS [73]. The activation of glial cells is the hallmark of inflammation in the brain [74]. Activated microglia produce neurotoxic inflammatory stress signals, such as, tumor necrosis factor-alpha (TNF-α) [28], interleukins (1β, IL-2, IL-6, IL-8 and IL-12) [75], lipoxygenase [76], cyclooxygenase-2 (COX-2) [77], monocyte chemoattractant protein (MCP) [26], growth factors and complement proteins [78]. These proinflammatory mediators, in turn, precipitate an inflammatory signaling cascade by activating other resident cells to produce additional molecules that perpetuates microglia activation in a positive feedback loop [73].

Increased inflammation due to chronic exposure to high fat diet increases vulnerability of neurotransmitter receptors to oxidative stress through activation of the oxidative stress-sensitive nuclear factor kappa-β (NF-kβ) [79]. The activated NF-kβ, in turn, upregulates the inflammatory response resulting in a further increase in ROS such as superoxide species and nitric oxide (NO) as well as increased expression of inducible nitric oxide synthase (iNOS) [80]. High levels of ROS exacerbate oxidative stress and inflammation, and thus, vulnerability to further stressors [81]. Facilitated central adiposity might also precipitate an oxidative damage due to compromised redox homeostatic status characterized by attenuated antioxidant defenses thus exacerbating a neuropsychiatric damage [30,82]. The therapeutic effects exhibited by the *G. glauca* leaf extract could be due to its ability to mitigate inflammation and oxidative stress by down-regulating the activity and release of proinflammatory mediators and restoration of redox homeostatic status through activation of antioxidant defenses [26]. The normalization of NF-kβ levels reduces the expression of the pro-inflammatory cytokines and consequently results in low levels of ROS in the hippocampus [83].

Pharmacological manipulations of anxiety disorder by anxiolytic agents like benzodiazepines (BDZs) and allo-pregnanolone enhances GABAergic tone. Binding of the anxiolytic agents to one of the two gamma sub-unit of the GABA-A receptor causes a structural modification of the receptor and allosterically increases GABA-A receptor activity [84,85]. This binding also facilitates opening of GABA activated chloride channels thereby increasing chloride ion conductance and inhibition of the action potential [86]. The eventual allosteric binding of GABA to the gamma sub-unit of the GABA-A receptor decreases the excitability of neurons and augments a calming effect [85,86]. The observed anxiolytic properties of the *G. glauca* leaf extract may be attributed to stimulation of the binding of gamma-aminobutyric acid (GABA) to GABA-A receptors that occurs abundantly on the surfaces of neurons in the amygdala and other parts of the limbic system and, therefore, brings a calming effect [25].

In the present study, the observed anxiolytic effects and increased locomotor and exploration-like behaviors in extract-treated rats could be attributed to the presence of a number of bioactive compounds in the DCM leaf extract of *G. glauca*. The synergistic and/or additive effects of these phytochemical compounds might be implicated in amelioration of symptomatic complications of obesity viz; anxiety, locomotor activity and exploration.

The phenolic compounds such as catechins and epicatechins were found to confer neuroprotective effects by mitigating oxidative and metabolic insults [87]. Catechins exhibit neuroprotective activities by activating multiple signaling pathways that exerts cell-survival and anti-inflammatory actions, including altering the expression of pro-apoptotic and anti-apoptotic proteins and upregulating antioxidant defenses [88]. Catechins activate protein kinase C (PKC) and transcription factors that promote the expression of cell-survival genes [89]. Studies reported that catechins and epicatechins exhibited protective effect of dopaminergic neurons from damage induced by 6-hydroxydopamine in a rat model of Parkinson’s disease [87]. In addition, catechin and epicatechin suppresses neuroinflammation, attenuate activation of microglia and inhibit release of the mediators associated with apoptotic death of neurons [90].

Curcumin was shown to ameliorate impaired hippocampal neurogenesis and increase the expression levels of brain-derived neurotrophic factor (BDNF) in severely stressed rats [62]. Curcumin has been used in the prevention and management of neurodegenerative diseases such as Alzheimer’s Disease, Parkinson’s Disease and stroke [62]. It mitigates oxidative stress and inflammation by down-regulating the activity of lipoxygenase, COX-2 and inhibiting the generation of proinflammatory cytokines such as TNF-α, interleukins I, II, VI, VIII and XII as well as monocyte chemoattractant protein (MCP) [75,91]. Besides, curcumin deactivate the transcription factor NF-kβ through induction of the expression of antioxidant enzymes and Bcl-2 [92]. Curcumin activate multiple signaling pathways through ligand binding to various receptors that include Growth Factor Receptors (GFR), G-Protein Coupled Receptors (GPCR) and Insulin Receptors (IR). These receptors in turn activate the kinase cascades involving Phosphatidylinositol-3-Kinase (PI3K), Mitogen-Activated Protein Kinases (MAPK) and Protein Kinase C (PKC) [92].

Quercetin has been shown to improve brain cell function and signaling by mitigating extra-neuronal parameters of survival—the oxidative stress [93]. Quercetin reduced oxidative stress and protected cultured hippocampal neurons against nitric-oxide-mediated cell death [87]. Quercetin ameliorates calcium dysregulation thereby protecting from ischemic injury, neuronal cell death and consequently brain damage [94]. Quercetin treatment decreased acid-mediated intracellular calcium levels and inhibited spectrin breakdown by inactivation of calcium-dependent protease cabin [95]. Quercetin significantly decreased protein oxidation, A*β*-induced toxicity and apoptosis in primary hippocampal cell cultures [94]. This novel antioxidant offers an effective and safe means of bolstering body’s defense against free radicals [96].

Stilbenes such as pinosylvin and resveratrol are phytophenols that have been shown to exhibit antioxidant activity [97]. Pinosylvin and resveratrol protected cultured neurons against oxidative damage by scavenging nitric oxide radicals [42]. Similarly, in a model relevant to Parkinson’s disease, resveratrol protected cultured dopaminergic neurons against oxidative-induced cell death [42]. Current findings indicate that administration of resveratrol and/or pinosylvin to rats confers protection of neurons in the brain and spinal cord from ischemic injury [42]. In models relevant to Alzheimer’s disease, stilbenes promoted clearance of amyloid β-peptide from cultured cells hence preventing neuronal cell damage [98].

Flavonoids were reported to modulate neuronal function and prevent neurodegeneration [99]. Flavonoids were shown to improve memory and learning through stimulation of neuronal regeneration and enhancement of neuronal function [92]. They inhibit TNF-α, IL-1β and nitric oxide in activated microglia cells [92]. Flavanone maintain nigro-striatal integrity and functionality and serve as a potential neuroprotective agent against 6-hydroxydopamine [99]. Flavonoids activate P13 kinase-mTOR cascade and ERE-CREB pathway resulting in changes in synaptic plasticity. Flavanones were found to inhibit oxidative-induced neuronal apoptosis through phosphorylation of signaling proteins essential in pro-survival pathways. Capsaicin have been observed to activate specific Ca^2+^ channels by its action on vanilloid receptors [92].

Previous studies reported that condensed tannins such as gallocatechin-catechin flavan and anthocyanins have the ability to diffuse through the central nervous system and cross the Blood-Brain Barrier (BBB) [100]. Gallocatechin-catechin flavan and anthocyanins confers neuroprotective function through their antioxidative properties. In human SH-SY5Y neuroblastoma cells, condensed tannins reduced A*β*-induced neurotoxicity by enhancing the formation of A*β* fibril formation thus reciprocally modulating local A*β* clearance [101]. Gallocatechin-catechin flavan and anthocyanins have been shown to have potent anti-inflammatory activities. They inhibit inflammatory mediators COX-2 [102].

Alkaloids increases gamma aminobutyric acid (GABA) in the synapses of the brain [103]. They are a highly potent vasodilator agents, enhances cerebral blood flow, facilitate glucose uptake by brain cells and protects from hypoxia and ischemia [103].

Ellison and α-amyrin were shown to activate the Transient Receptor Potential (TRP) ion channels in the cell membrane of neurons [104]. This resulted in Ca^2+^ influx which in turn activates neuroprotective kinase signaling cascades via Camp-Response-Element-Binding protein (CREB) and Mitogen-Activated Protein Kinases [104]. The CREB stimulates expression levels of a major neurotrophic factor, the Brain Derived Neurotrophic Factor (BDNF). The BDNF activates the PI-3K/Akt and MAPK/ ERK pathways through binding to its tyrosine kinase TrkB receptor thereby activating downstream molecules which can promote neurogenesis and cell survival [105].

Ferulic acids attenuates the stress-induced behavior in the depression-like model in mice [101]. Ferulic acids influence the function of ionotropic receptors for GABA in the brain, therefore, enhancing its anxiolytic effects [92].

Terpenoids (such as monoterpenes, triterpenoid and sesquiterpene alkaloid) isolated from the rhizome of *Valerian officinalis* exhibited a broad range of neuroprotective actions [106]. Terpenoids were shown to confer sedating effects in mice through the activation of GABA-A receptor activity and other pathways upstream of nuclear factor erythroid 2-related factor 2 (Nrf2) [106].

Polyunsaturated fatty acids (PUFAs) such as oleic acid, linoleic acid and α-linolenic acid maintain the integrity of the structural components of neurons [107]. The fatty acid composition of the neuronal membrane is necessary for maintenance of appropriate electrical gradients across the membrane and neurotransmission in the synaptic cleft [108]. The PUFAs improve membrane fluidity and, therefore, affects membrane biophysical properties. In neuronal membranes, PUFAs participates in signaling cascades that promote synaptic plasticity, neuronal function and neuroprotection [109].

## 1.5 Conclusion

Chronic consumption of a high-fat diet has been implicated in impairments of neurobehavioral domains such as locomotor activity, anxiety and exploration. The present study demonstrated that a higher frequency of cross lattice, rearing and center square entries are indicative of increased locomotion and exploration-like behaviors. The high frequency of fecal, urination and stretch attend postures score are indicative of anxiety. Besides, the increased freezing period, grooming patterns and latency duration are suggestive of increased anxious states. The therapeutic application of graded doses of the *G. glauca* indicated anxiolytic effects and increased spontaneous locomotor activity and exploration-like behaviors in HFD-induced obese rats. The observed therapeutic effects might be attributed to the phytochemicals contained in the *G. glauca*. These phytocompounds have been shown to mitigate the main mediators of neurodegeneration viz inflammation and oxidative stress by down-regulating the activity and release of proinflammatory mediators and restoration of redox homeostatic status through activation of antioxidant defenses. The findings of the present study provide ‘qualified leads’ for the synthesis of new alternative antioxidant suppliment and therapeutic agent for management of obesity and other associated symptomatic complications such as anxiety and impaired motor faculties. However, there’s need for further studies to establish the depth of this possibility.

## Acknowledgements

The authors wish to acknowledge the Department of Biochemistry, Microbiology and Biotechnology of Kenyatta University for allowing us to use the departmental animal house facility for rats breeding and performing the efficacy studies. Authors also acknowledges Mr. Joshua Mulele Machayo of the Department of Physical Science (Biochemistry Section) of Chuka University and, Mr. Daniel Gitonga Mwaniki and Joyce of the Department of Biochemistry, Microbiology and Biotechnology of Kenyatta University for technical support.

## Authors Contributions

Arika Wycliffe Makori carried out the study and wrote the manuscript. Dr. Cromwell Kibiti, Dr. Joan Murugi and Dr. Mathew Piero Ngugi contributed to conception of the review and supervised the manuscript writing. All authors have read and approved the final manuscript.

## Ethical Approval

The experimental protocols and procedures used in this study were approved by the Ethics Committee for the Care and Use of Laboratory Animals of Kenyatta University, Kenya.

## Data Availability

No data was used to support this study.

